# Molecular insights into conformational dynamics associated with the open-closed-phosphorylated states of PITPα

**DOI:** 10.1101/152587

**Authors:** Chaithanya Kotyada

## Abstract

Phosphatidylinositol transfer proteins (PITPα) are lipid carrier proteins that are involved in replenishing the phosphatidylinositol lipid molecules on the plasma membranes. Transitions between the open and closed state conformations are necessary steps in the mechanism of lipid transfer by PITPα. The apo (open state) conformation is assumed to occur at the membrane surface during the lipid exchange while the lipid bound (closed state) conformation is required for transfer activity. The transfer of phosphatidylinositol by PITPα is controlled by the phosphorylation of S166 in its regulatory region. We wanted to decipher the molecular basis for structure-function relationship between the open-closed-phosphorylated states of PITPα. We used all-atom Molecular Dynamics Simulations to study the conformational dynamics in each of these states. Our study shows that the open state is highly dynamic and its transition to closed state would stabilize PITPα. We observed restricted conformational sampling of the phosphorylated state which provide basis for its decreased lipid transfer activity. Further, using analysis of residue-residue contact maps and hydrogen bond interactions we discuss the impact of phosphorylation on the global conformation of PITPα. Overall, our work provides insights into the structural dynamics in each state and their functional significance.

## Introduction

Phosphatidylinositol signalling plays a key role in many signal transduction pathways in the cell. The PI on the plasma membrane is phosphorylated to PI(4,5)P2 which is an important substrate for PLC mediated signalling[1-3]. The phosphatidylinositol (PI) levels needs to be maintained on the plasma membrane for normal cellular signalling. Transfer of newly synthesized lipids to the membrane can be achieved by two mechanisms (1) vesicular trafficking between membrane compartments, (2) Lipid carrier proteins mediated transfer of the newly synthesized lipids onto the destination membrane compartment[4,5]. Phosphatidylinositol transfer proteins (PITPs) are lipid carrier proteins that balance the membrane lipid concentration by shuttling the phospholipids between the compartments. The major function of PITP (α, β) is to maintain the phosphoinositide levels by specifically exchanging phosphatidylinositol (PI) for phosphatidylcholine (PC) and vice-versa[5]. The transfer activity is regulated by phosphorylation of PITP by PKC[5-7]. Phosphorylation of PITPα at S166 residue attenuates the PI transfer ability of the protein[7].

Crystal structures are available for the apo, PC and PI bound states of PITPα[8-10]. Comparing the crystal structures show two predominant conformations for the apo (open state) and the lipid bound (closed state). In the closed state the lipid molecule is completely shielded from the surrounding aqueous environment (Fig. 1A). The two states show major conformational deviations in the G-helix, C-terminal tail and the lipid exchange loop (Fig. 1A). Biophysical studies confirm the major transitions in the G-helix and lipid exchange loop[11,12]. Similarly, mutational analysis indicate active role for the W202/W203 residues in the membrane recognition of PITPα[13]. The lipid transfer mechanism involves several steps in the following order (1) membrane recognition (2) closed-open state transition (3) exchange of bound lipid (4) open-closed state transition and (5) removal of the protein from the membrane. Studies were done previously on similar systems to comprehensively understand the mechanistic details of each of these steps[14,15]. Though, MD simulation studies on the functionally similar Sec14 protein from yeast revealed some details regarding the gating mechanism, the sequence-structure variations between two proteins is very large to derive any correlation[16-18]. The mechanistic details regarding the lipid transfer activity by PITPs largely remains unexplored.

**Figure 1.**
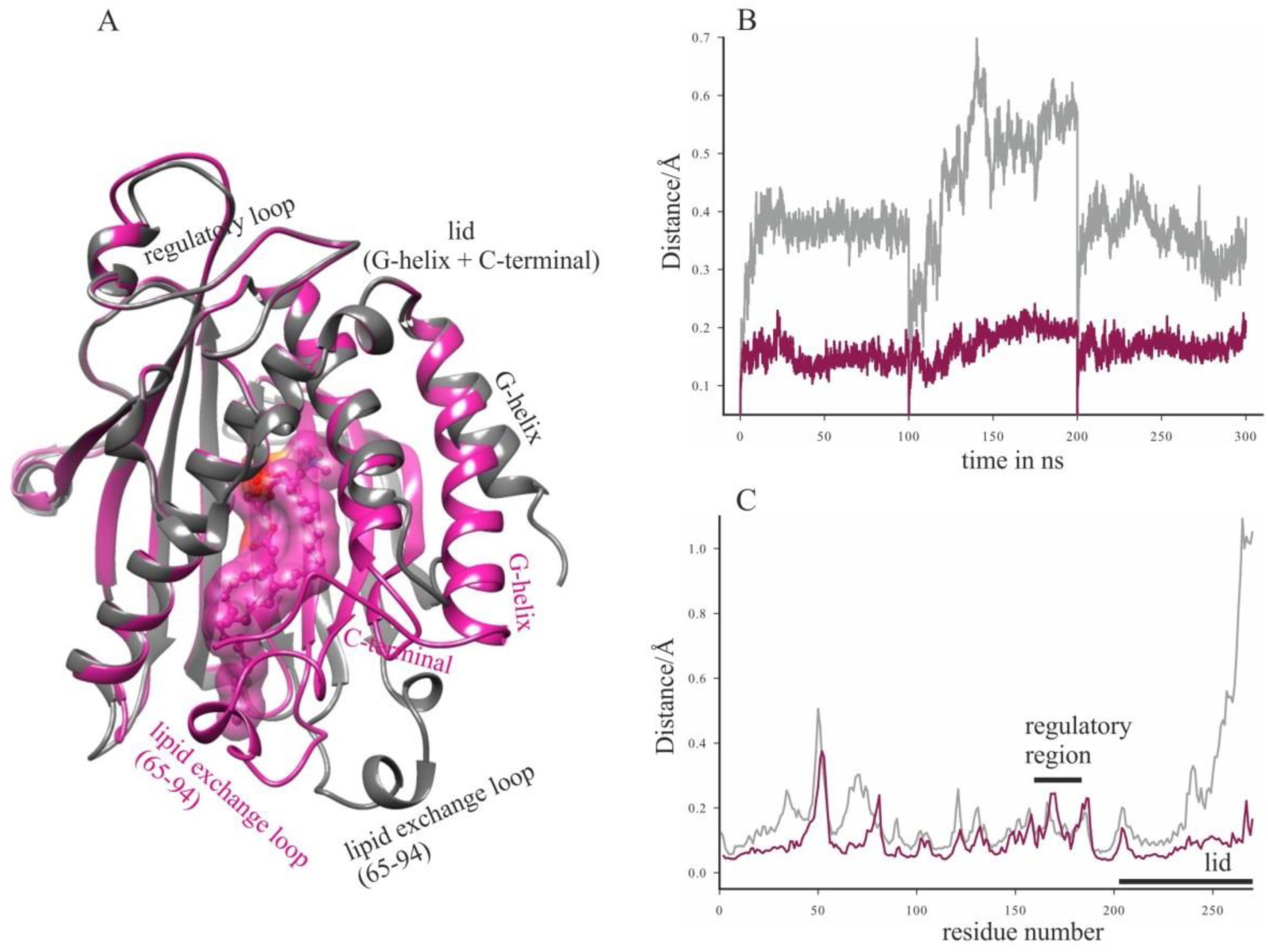
Comparision of open and closed states of PITPα. (A) Overlay of crystal structures of open PITPα (PDB ID 1KCM) dark grey, closed PITPα (PBD ID 1T27) violet red, representing the conformational variation. (B) Overlay of RMSD plots of the open PITPα (dark grey) and closed PITPα (violet red). (C) Overlay of RMSF plots of the open PITPα (dark grey) and closed PITPα (violet red), representing C-α fluctuations at the lid and regulatory regions.

The open, closed and phosphorylated states of PITPα are functional states that regulate its activity in the cell. The abrogation of functional activity of PITP is associated with many cancers and neurological disorders[19]. We wanted to understand the molecular basis for the functional dynamics of each of these states. We used all-atom MD simulations to study the conformational dynamics associated with each of these states. Our analysis on the 6000 conformations obtained for each state reveal the structural heterogeneity between all the three states. We provide molecular basis for efficient transition from open state to closed states. Further, we detail the basis for inhibition of PI transfer by the phosphorylated state of PITPα. PITP’s are also found be domains in larger proteins like retinal degeneration (RdgB) proteins [20,21]. Therefore, our study provides preliminary insights into the mechanistic basis for the lipid transfer activity of PITP like proteins.

## Methods

### Model Preparation

For the MD simulations of open and closed state, initial models were prepared from the crystal structures of apo PITPα (PDB ID 1KCM) and phosphotidylcholine bound PITPα (PDB ID 1T27) respectively. The closed state model is used as template to generate the double charged phosphoserine (SP166) residue at the S166 position using the swisssidechain plugin of UCSF Chimera[22]. Crystallographic water molecules were removed from all the models prior to system preparation.

### Molecular Dynamic Simulations

All-atom MD simulations were performed using GROMACS simulation package, version 5.0.4. The system was parameterized using CHARMM36 forcefield and TIP3P rigid water model. The parameters of the single lipid of DOPC membrane in CHRAMM36 forcefield was used for the phosphotiydlcholine lipid in the protein-lipid system. The initial models were placed in a cubic box with the boundary at least 10Å from the boundary of the system and TIP3P rigid water. Appropriate counter ions were added to neutralize the total charge of the system. Steepest descent algorithm was used to energy minimize the system until the system converged with Fmax no greater than 1000 kJ mol−1 nm−1. Equilibrations was performed under NVT conditions for 600ps and NPT conditions for 1200ns. The v-rescale and Berendsen thermostats were used to couple temperature at 300K and pressure 1bar. LINCS algorithm was used to constrain all bond angles and the electrostatic interactions were evaluated using Particle Mesh Ewald (PME) method. Three independent trajectories of 100ns were then performed on the equilibrated open, closed and phosphorylated systems using leap-frog algorithm.

### Analysis of trajectories

Analysis was performed on the 6000 conformations generated for the entire 300ns MD run for each system. RMSD, RMSF, PCA and SASA calculations were performed using GROMACS tools. Timeline plugin of VMD was used for secondary structure analysis of the systems. Percentage of hydrogen bond occupancy measurements between residues was done using the VMD tool. UCSF Chimera software package was used for measurement of distance between two atoms, cluster analysis and generation of final images.

## Results

### Open to closed state transition is accompanied by drastic reduction in the molecular fluctuations of PITPα

Transition between the open (apo) and closed (lipid bound) state conformations is an essential step in the phosphatidylinositol transfer activity of PITPα. Crystal structures of the open (PDB ID 4KCM) and closed (PDB ID 1T27) states show differences in the C-terminal (G-helix and the downstream tail residues) designated as lid and the residues corresponding to 65-94, referred to as the lipid exchange loop (Fig. 1A). Understanding the conformational dynamics associated with these states may provide insights into the mechanistic basis for open to closed state transition. We used all-atom molecular dynamics simulations to study the conformational dynamics in each of these states. Initial models for the open and closed states were generated from the crystal structures of apo (PDB ID 1KCM) and PC bound PITPα (PDB ID 1T27), respectively. Three independent trajectories of 100ns each were performed on the initial models of open and closed states. Figure 1B. shows the root mean square deviations (RMSD) of backbone atoms for the 300ns production run of the open state and closed state conformations of PTIPα. The backbone RMSD values are very high in the open state compared to the closed state (Fig. 1B). The deviations reaches a maximum of 4-6A for the open state and 1-2A for closed state (Fig. 1B). To identify the regions contributing to the large deviations in the open state, we plotted the root mean square fluctuations (RMSF) of the backbone C-α atoms in the open and closed states of the protein (Fig. 1C). The open state shows overall increase in the C-α fluctuations throughout the protein (Fig. 1C). However, maximum fluctuations were observed in the lid region (Fig. 1C). Interestingly, the RMSF of the regulatory loop seems to be similar in both open and closed states (Fig. 1C). These results indicate that the binding of PC to PITPα stabilizes the protein by drastically reducing its molecular fluctuations. The huge reduction in the dynamic flexibility would facilitate efficient transition from the open state to more stable close state of PITPα.

### Phosphorylation of Ser166 confines the conformational sampling of PITPα

Previous studies showed that the S166 residue in the regulatory loop of PITPα is phosphorylated by PKC[7]. Phosphorylation of S166 hinders the PI transfer activity of PITPα, despite retaining its ability to bind to the lipid[6,7]. The lipid bound conformations of PITPα is known to be good substrates for PKC (ref). We wanted to understand the molecular basis governing the reduced transfer activity of the protein. We used the closed state as template to generate in-silico phosphoserine (SP166) at the S166 position. Simulations were performed for a total of 300ns with three independent trajectories of 100ns each. The replacement of S166 by SP166 did not affect the secondary structure of the protein(Sppl. Fig. 1). The RMSD values also remained similar for both the closed state and phosphorylated state (Sppl. Fig. 2). However, the C-α RMSF is not consistent between the two states (Fig. 2). The phosphorylated state becomes flexible at the regions corresponding to helix 205-230, N-terand the regulatory loop. Interestingly, the helix 205-230 borders the lipid tail originating fromsn2 carbon of PC(Fig. 2). The closed state shows maximum deviation from the phosphorylated state at the A81 of the lipid exchange loop(Fig. 2). We plotted φ-ψ scattered plots for the residues at the interaction surface of the lipid exchange loop (Fig. 3). Ramachandran plots revealed restricted φ-ψ distribution for the residues L65, A81, L82, Y103, W202 in the phosphorylated state (Fig. 3). Previously, W202 andW203are shown to be critical for the membrane recognition by PITPα[13]. Similarly, the side-chain conformation of Y103 was shown to cap the sn2 lipid tail of phospholipid[5]. Mutations of these residues are shown to either fully or partially compromise the binding-transfer activity of PITPα[8,13]. We hypothesized that the phosphorylation of PITPα may be restricting the global conformational sampling of PITPα. We performed cluster analysis for the 6000 frames of closed and phosphorylated states of PITPα with a step size of 50 frames. In the closed state the conformations were distributed into 14 cluster ensembles compared to 4 cluster ensembles of phosphorylated state (Fig. 4). Similarly, the cluster ensembles for the PC were 19 and 9 for the closed and phosphorylated states, respectively (Sppl. Fig. 3A). The occasional bending of the sn1 lipid tail towards the lid region in the closed state is noteworthy (Sppl. Fig. 3B). Evidently, both the protein and lipid showed defined ensemble of clusters in the phosphorylated state compared to the closed state. Therefore, the phosphorylation of S166 residue in PITPα restricts the conformational sampling of both protein and lipid.

**Figure 2.**
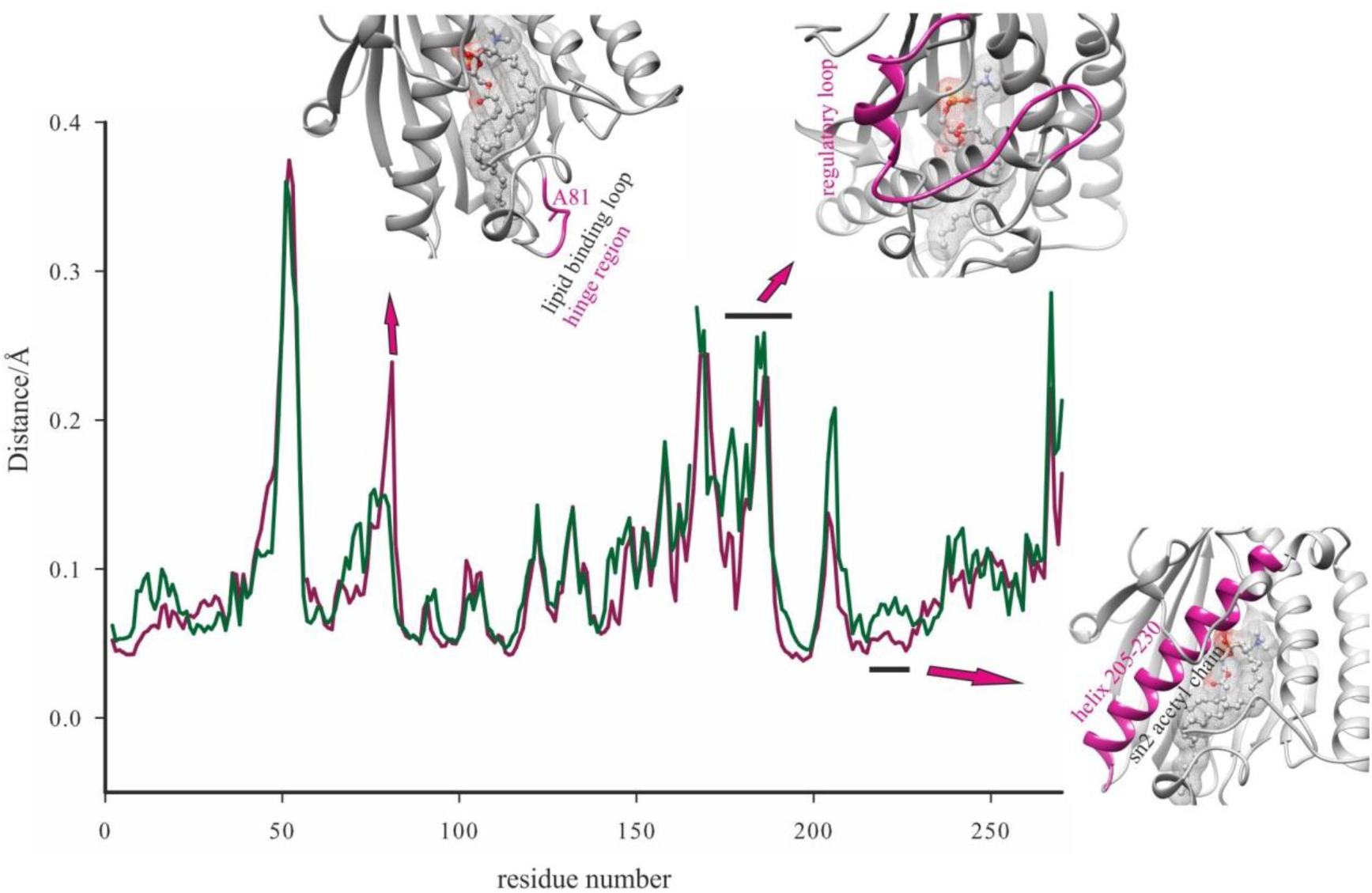
Overlay of RMSF plots of the closed PITPα (violet red) and phsophorylated PITPα (dark green). Cartoon representation of the flexible C-a regions in the phosphorylated state.

**Figure 3.**
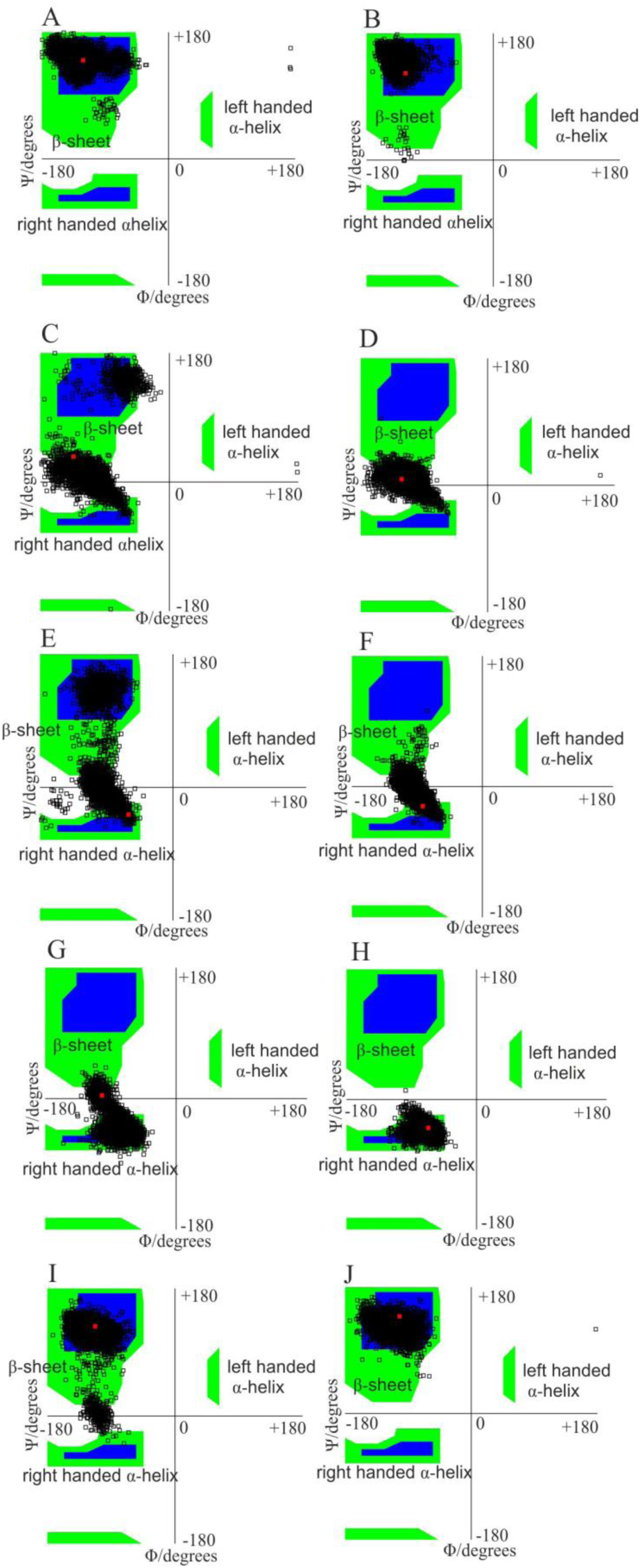
Ramachandran Plots. Left are the Φ-ψ distributions for the residues (A) L65, (C) A81, (E) L82, (G) Y103, (I) W202 in the closed state, Right are the Φ-ψ distributions for the residues (B) L65, (D) A81, (F) L82, (H) Y103, (J) W202 in the phosphorylated state.

**Figure 4.**
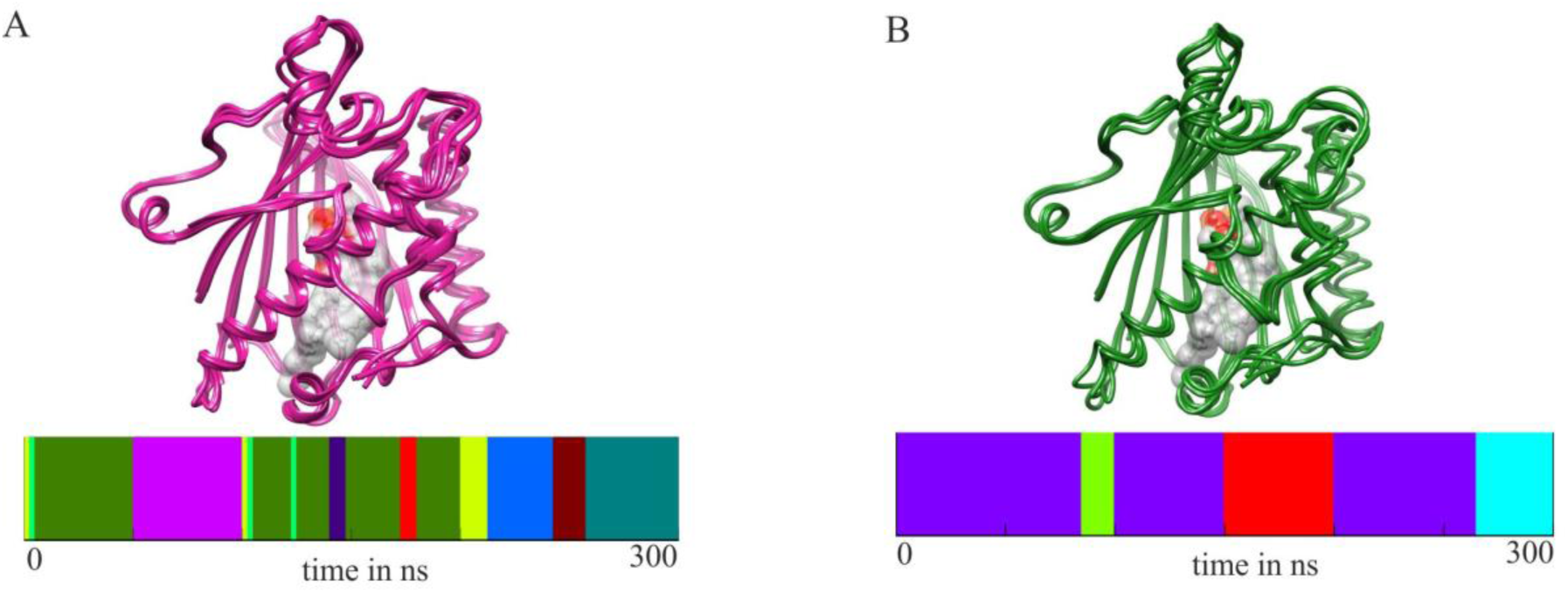
Cluster Analysis. (A) Overlay of the average structures of the four major ensembles of the fourteen in the 300ns simulation of closed state (violet red, above), Distribution of the ensemble conformations over the 300ns simulations (below). (B) Overlay of the average structures of the all the four ensembles in the 300ns simulation of phosphorylated state (dark green, above), Distribution of the ensemble conformations over the 300ns simulations (below).

### Notable changes in the global residue interactions and water dynamics due to phosphorylation of S166 in PITPα

To understand whether phosphorylation induces any global change in the residue interaction network, we calculated the residue-residue contact maps for both the closed and phosphorylated states of PITPα. Interestingly, some of the residue-residue contacts in the closed state disappear upon phosphorylation (Fig. 5). We then calculated the overall hydrogen bonds in the closed and phosphorylated states. Upon phosphorylation of S166 residue the hydrogen bond interaction between D234-S166, T169-S166 is replaced by D234-T169 (Fig. 6A). This local change in the interaction showed consequences on the global hydrogen bond interaction pattern. The percentage occupancy of hydrogen bonds between residue pairs varied among the closed and phosphorylated states. Notable changes in the hydrogen bond interactions include K195-T114, T59-N45 (Fig. 6B). The residues K195, T114,Q22 are involved in direct and water mediated interactions with the oxygen atoms of phosphate, respectively (Fig. 6B). We suggest phosphorylation induces notable changes in the residue interaction pattern of PITPα.

**Figure 5.**
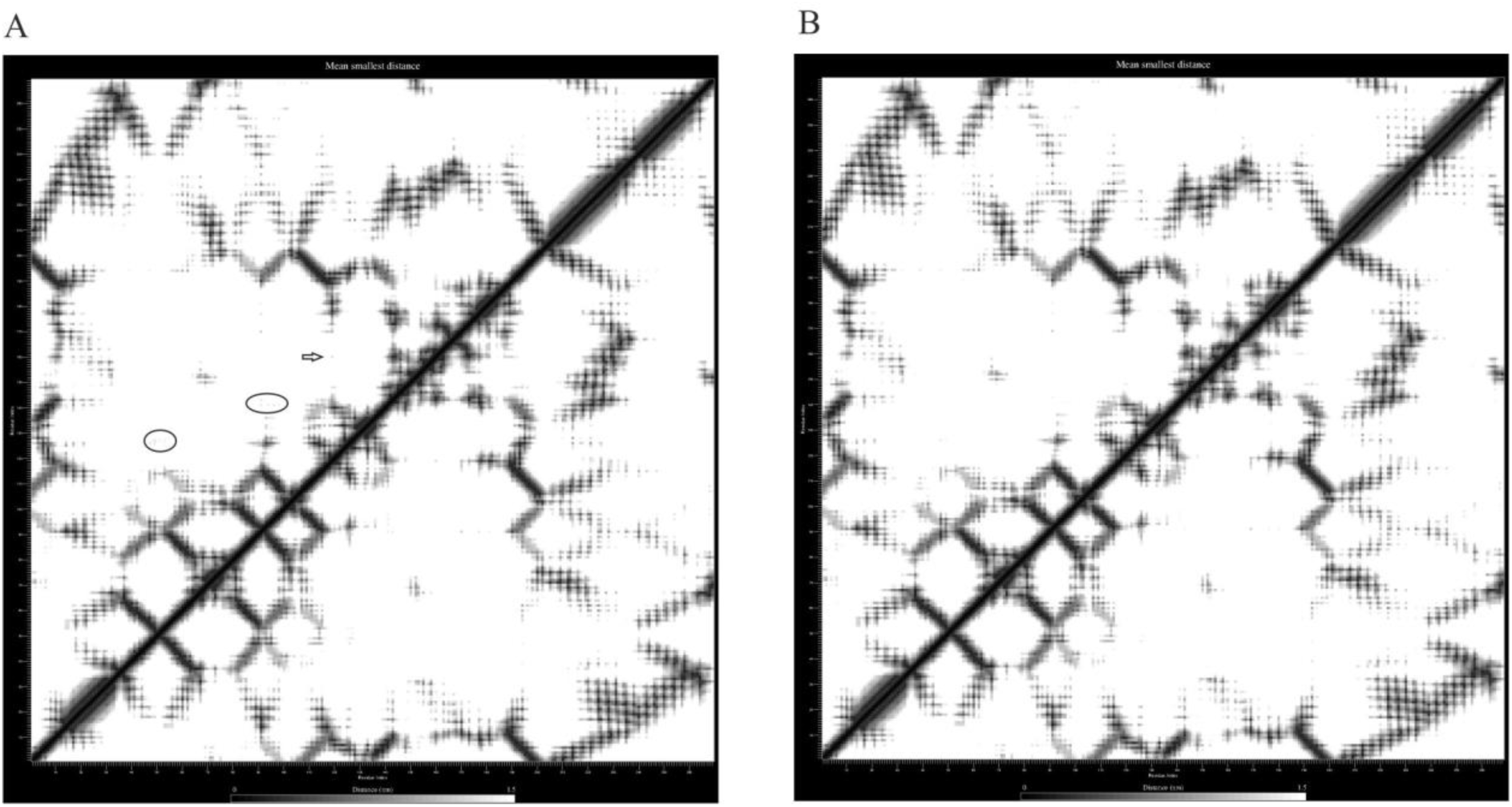
Residue-residue contact map. (A) closed state, highlighted residue interactions disappear upon phosphorylation. (B) phosphorylated state.

**Figure 6.**
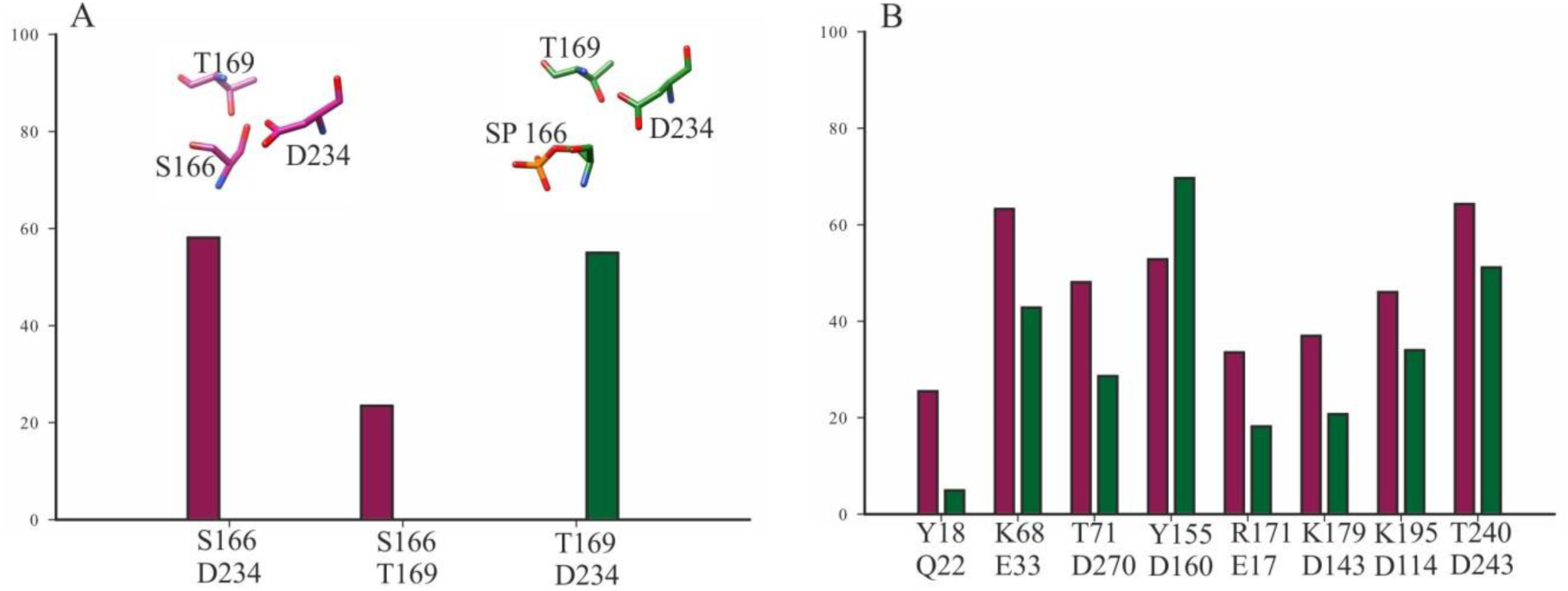
Hydrogen bond occupancy measurements. (A) Hydrogen bond interactions at the phosphorylated site S166, before phosphorylation (violet red), after phosphorylation (dark green). (B) Comparison of hydrogen bond interactions in the closed (violet red) and phosphorylated state (dark green).

We performed surface area solvent accessibility (SASA) measurements for C-αatoms in the open and closed states of PITPα. Interestingly, the SASA measurement is similar for both the states except at the regulatory region (Fig. 7A). The S166 residue is buried in the regulatory loop and is not accessible for phosphorylation by PKC in the closed state (Fig. 7A). Studies assume that alternate conformation of regulatory region is required for efficient phosphorylation of PITPα[6]. The increased SASA of the regulatory region indeed concurs with the necessity for distinct PITPα conformation as substrate for PKC. We next analyzed the dynamics of water molecules interacting with the residues of PITPα. T114 and Q22 are involved in water mediated interactions with oxygen atom of phosphate group of lipid (Fig. 7B). Interestingly, the residence time for the T114 bridging water molecule increases upon phosphorylation (Fig. 7B). These results suggest differential water dynamics in the closed and phosphorylated states of PITPα.

### PCA show distinct collective motions for the open, closed and phosphorylated states of PITPα

To evaluate the differences in the correlated dynamics of the three states, we performed principal component analysis of backbone atoms on the MD simulation trajectories of each system. The projections of PC1 and PC2 for backbone atoms of each system were plotted as abscissa and ordinate, respectively. The conformational sampling of the open state is highly diffused with three minor cluster ensembles (Fig. 8A). In contrast the closed and phosphorylated states show localized collective motions (Fig. 7B-7D). The PCA of closed state is distributed into three cluster ensembles (cluster1, cluster2, cluster3) compared to the two distinct cluster ensembles (cluster1 and cluster2) of the phosphorylated state (Fig. 8B-7D). Evidently, the restricted collective motions in the phosphorylated state suggest that protein adopts limited conformations compared to the closed state. Clearly, lipid binding facilitates the open to close state transition by drastically reducing the collective dynamics of the protein. Further, phosphorylation of S166 residueconstrains the functionally relevant motions to distinct clusters.

**Figure 7.**
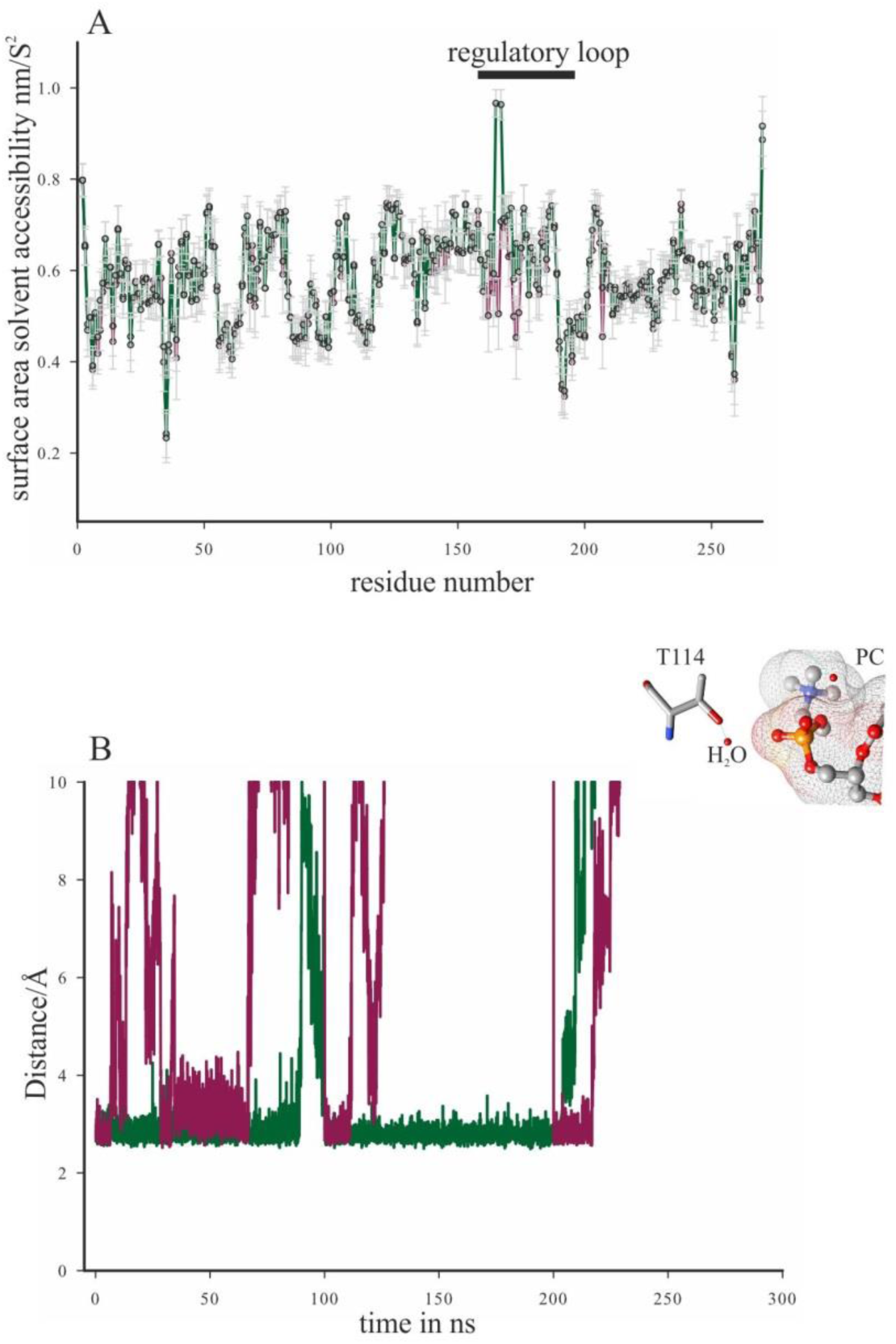
Difference between the water dynamics of closed and phosphorylated states. (A) Surface area solvent accessibility measurements (SASA) in the closed state (violet red), and phosphorylated state (dark green). (B) Distance measurements between the oxygen atom of bridging water molecule and phosphate of phospholipid over the 300ns simulation time, closed state (violet red), phosphorylated state (dark green).

**Figure 8.**
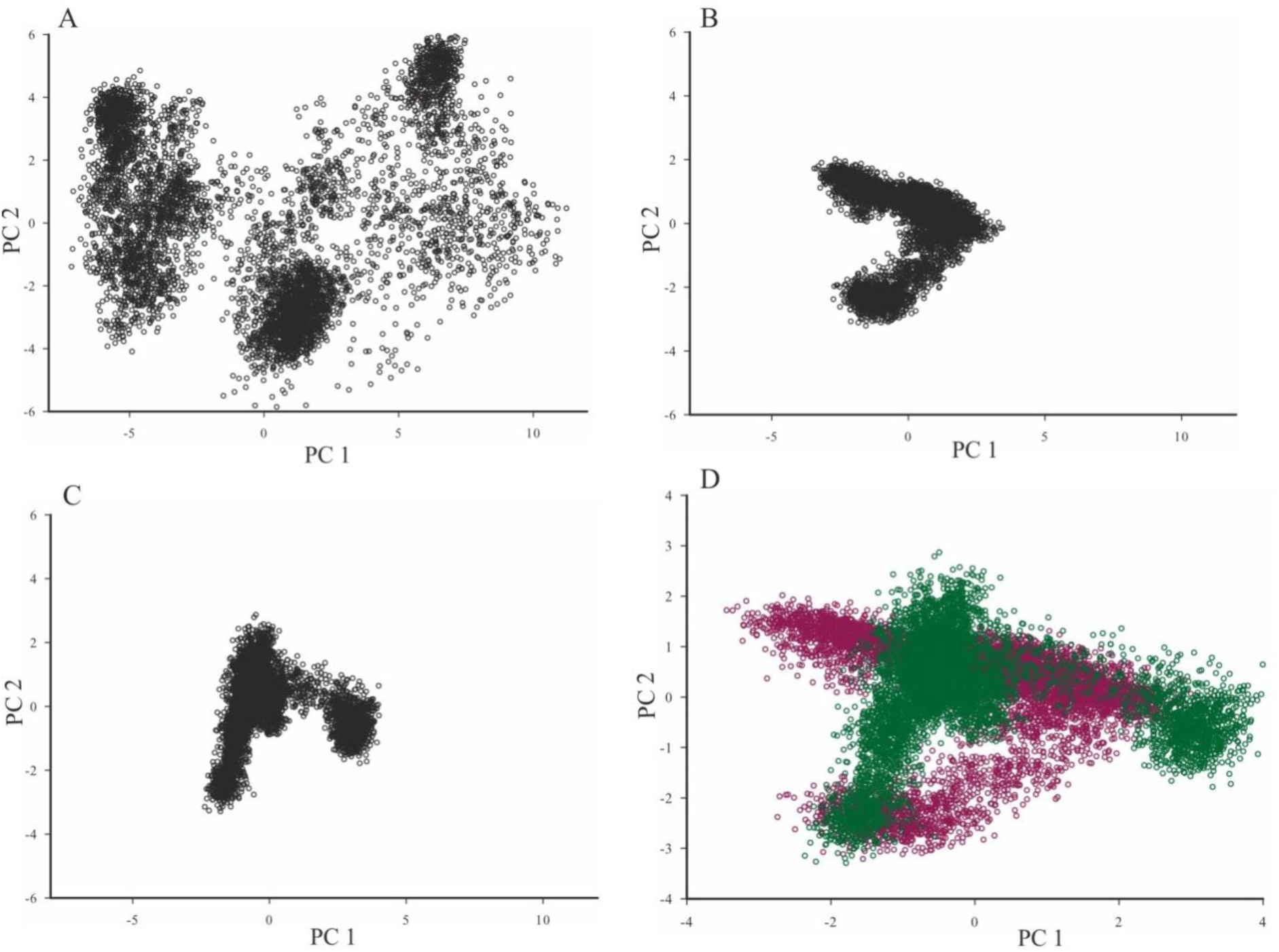
Principal Componant Analysis of PITPα. (A) open state (B) closed state (C) phosphorylated state (D) overlay of closed and phosphorylated states.

## Discussion

In this study we describe the differences observed in the dynamics of PITPα in its open, closed and phosphorylated states. We show that the high intrinsic molecular fluctuations would make the open state very unstable. Sec14 in yeast is a functional analogous protein of PITPα which possess a conformational gating mechanism for the open and closed states[16,17]. Sec14 does not share either sequence or conformational similarity with PITPα[ 16]. The gating mechanism in Sec14 involves opening and closing of a helix region[16]. Unlike, Sec14 the open and closed states shows major differences in the G-helix including C-terminal (lid) and short helical region spanning residues 65-94, which we designate as cap. The proposed mechanism of PI transfer by PITPαinvolves transient open state formation at the membrane region during lipid exchange[13]. We suggest that the open state achieves transient stability due to the protein-membrane interactions during membrane recognition. The immediate entry of phosphoplipidinto the pocketof PITPαfurther provides the momentum for efficient transition to the closed state.

The main function of PITPα protein is to transfer newly synthesized PI from endoplasmic reticulum to plasma membrane[23,24]. The transfer activity is controlled by the phosphorylation of PITPα by PKC[7]. Cellular studies show that the phosphorylation of S166 reduces re-localization of PITPα to perinuclear golgi structures[7]. The phosphomimmetic S166E is shown to severely impair lipid binding and transfer activity[6]. We wanted to understand the molecular basis for diminished membrane localization and transfer activity by the phosphorylated PITPα. Our observations indicate that the phosphorylation of PITPα restricts its conformational sampling. The PCA and structural ensemble analysis show that the conformational heterogeneity of phosphorylated state is concentrated to specific ensembles compared to the closed state (Fig. 8). We propose that the phosphorylated state of PITPα does not sample the conformations relevant for membrane recognition and lipid exchange. Therefore, phosphorylation would severely effect lipid binding, membrane recognition and lipid transfer.

The crystal structure of lipid bound PITPα shows that S166 residue is not accessible for phosphorylation by PKC. Studies assume that PITPα needs to adopt conformation in which S166 is exposed for efficient phosphorylation by PKC[6]. The way PITPα achieves PKC substrate specific conformation, is still not clear. However, the T59A mutant of PITPα is shown to be effective substrate compared to the wild-type[6].We see that the regulatory region of phosphorylated state is more accessible to the solvent water compared to the closed state (Fig. 7A).In addition, the phosphorylated state also shows differences in the hydrogen bonds at several positions in PITPα. These results show that the phosphorylation of S166 induces changes in the water dynamics of PITPα.

Taken together, our observations provide first step in understanding the mechanistic basis for lipid transfer activity by PITPα. Our study provides molecular basis for varying functional activity of the open, closed and phosphorylated states of PITPα. Further, studies needs to be carried out to arrive at specific details of membrane recognition and lipid exchange by PITPα

